# Expansion of Plant Redox Protein Network Predated Plant Terrestrialization

**DOI:** 10.64898/2026.05.07.723422

**Authors:** Ran Ovadia, Einat Hazkani-Covo, Shilo Rosenwasser

## Abstract

The evolutionary transition of the green plant lineage (*Viridiplantae*) from aquatic environments to terrestrial habitats required unprecedented adaptations of cellular metabolism to severe environmental stressors, including desiccation, high irradiance, and rapid temperature fluctuations. Redox regulation, mediated by oxidation and reduction of reactive cysteine residues (RCys), plays a crucial role in translating environmental fluctuations into rapid cellular responses. Although comparative genomics has revealed expansions in multiple cellular systems preceding terrestrialization, the evolutionary history of redox-regulated protein networks remains elusive. This work integrated large-scale phylogenomic reconstructions across 37 *Viridiplantae* species with five independent *Arabidopsis thaliana* redox proteomics datasets to trace the evolutionary trajectory of RCys. The analysis showed that the ancestral core, consisting of plastid-localized regulatory cysteines, was already established at the base of the green lineage. Furthermore, an expansion driven by gains of RCys via amino acid replacements within pre-existing proteins occurred in the common ancestor of Zygnematophyceae and land plants. These findings suggest that a targeted incorporation of thiol-based regulatory switches provided early land plant ancestors with enhanced protein functional plasticity necessary to cope with the challenges of terrestrial environments.

**Highlights:**

- The foundational plastid-localized redox core was established at the root of *Viridiplantae*.
- Novel regulatory switches were integrated into conserved machinery via amino acid replacement.
- A punctuated burst of redox innovation at Zygnematophyceae and Embryophyta last common ancestor preceded plant terrestrialization.
- Redox acquisition rates declined sharply following the successful colonization of land.

## Results and Discussion

The first photosynthetic eukaryotes emerged approximately two billion years ago through the primary endosymbiotic incorporation of a cyanobacterial progenitor into a eukaryotic host cell ^1,2^. This foundational event was followed by the diversification of the green lineage (Chloroplastida), roughly one billion years ago, which ultimately culminated in the colonization of terrestrial environments around 550 million years ago ^3^. Notably, the evolution of land plants (Embryophyta) can be traced back to streptophyte algal progenitors, representing a singular evolutionary transition rather than multiple independent origins ^4^. This momentous colonization of land marked a pivotal shift in Earth’s history, driving the rise of complex photosynthetic organisms and profoundly reshaping the biosphere, including global biogeochemical, carbon, and oxygen cycles ^5^.

A fundamental question in plant evolution concerns the traits that enabled the ancestral lineage to colonize terrestrial environments. Current phylogenomic and fossil evidence indicate that this transition was driven by a suite of preadaptations that had already evolved within streptophyte algae prior to the water-to-land transition ^6–12^. These included structural innovations, such as multicellularity and rhizoid-based anchoring systems, along with the emergence of advanced stress-signaling pathways and hormone-regulated networks critical for survival in the terrestrial niche.

Beyond these well-characterized structural and genetic adaptations, successful terrestrialization also depended on the ability to dynamically regulate metabolism in response to highly variable and harsh environmental conditions, a trait that in modern plants is achieved through redox-regulated protein networks ^13,14^. However, the evolutionary timing of these redox regulatory system acquisitions across the green lineage remains largely unexplored.

### Phylogenomic Framework for Redox Evolution

To reconstruct the evolutionary history of plant redox regulation, a high-resolution cysteine reactivity map for *Arabidopsis thaliana* was established by integrating data from five independent *in vivo* redox proteomics studies (Table S1). Integrating multiple datasets is essential to overcome the inherent coverage limitations of Liquid Chromatography-Mass Spectrometry (LC-MS)-based proteomics, as no single datasets comprehensively capture the redox-regulatory landscape. These datasets include a wide range of biochemical techniques and physiological conditions to identify redox-sensitive cysteines. Specifically, profiles of H_2_O_2_-induced oxidation in whole-cell lysates ^15^, cell cultures (Wei et al., 2020), and intact leaves ^17^ were incorporated. Data from seedlings exposed to excess light ^18^, and dark-to-light transition ^19^ were also integrated to capture light-dependent redox dynamics.

Cysteine sites were categorized into three distinct functional classes: **RCys** (reactive cysteines, based on study-specific reactivity thresholds), **NRCys** (non-reactive cysteines, detected by MS but falling below oxidation thresholds), and **UNCys** (unclassified cysteines, which reside in proteins where at least one other peptide was detected, but the specific Cys-containing peptide was missed). Overall, 967 RCys and 8077 NRCys sites were extracted. Analysis of the contribution of individual experiments to the compiled datasets revealed a minimal overlap. For example, out of 4,904 total sites (RCys and NRCys) detected in the largest single study ^17^, only 666 sites were consistently shared among the three largest datasets (Figure 1S). The filtered RCys landscape was even more exclusive; of the 640 RCys sites identified via Doron et al. ^17^, 608 were unique to that study. The largest shared subset consisted of only 14 RCys sites with a single subsequent study utilizing similar methodologies ^19^ (Figure 1A). This small overlap highlights the advantage of integrating data from independent studies to expand the repertoire of reactive cysteines.

**Figure 1.**
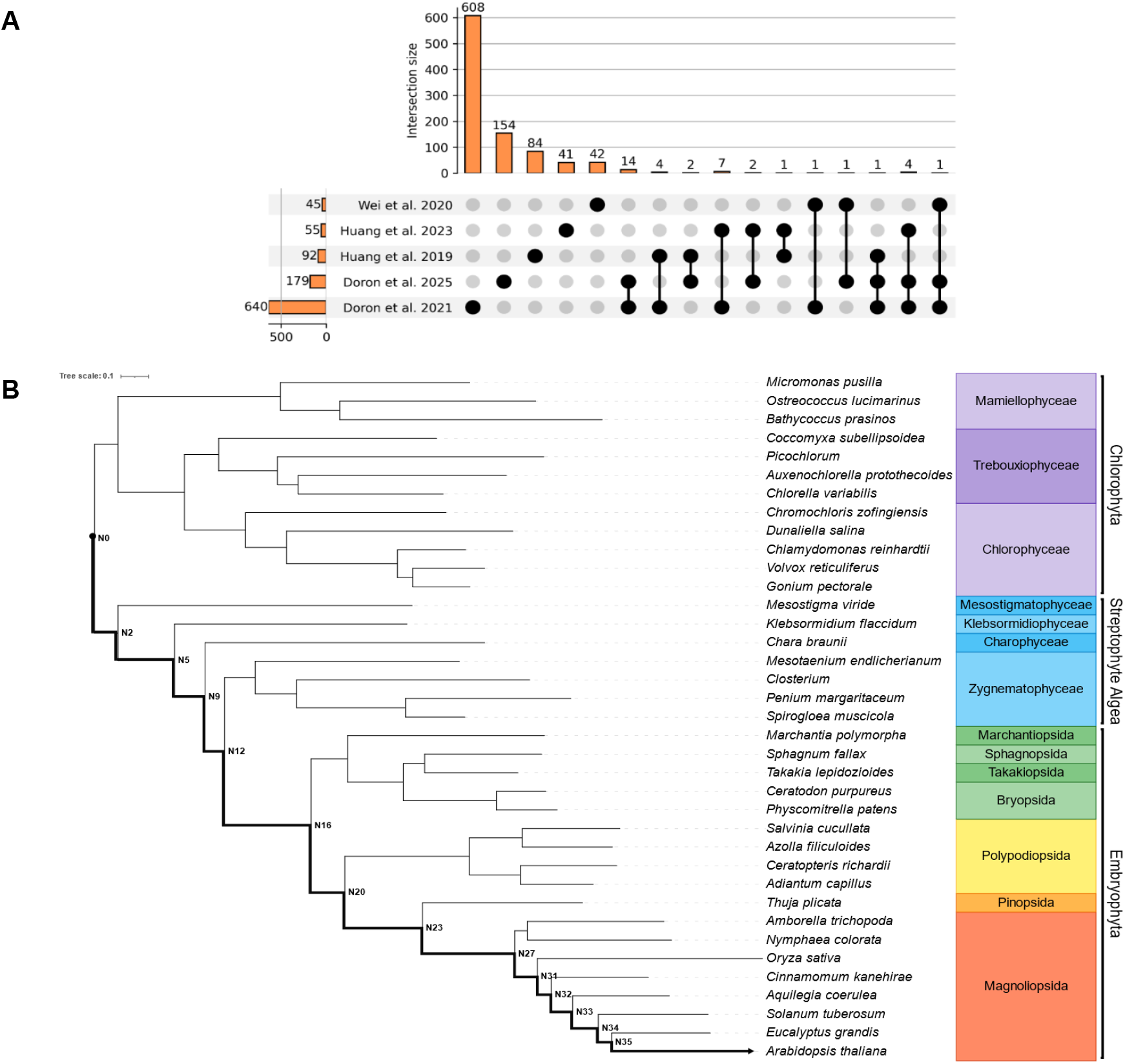
Integration of Redox Proteomics and Phylogenomics to Reconstruct the Evolutionary History of the Green Lineage. (A) Intersection of total cysteine sites across five independent *Arabidopsis* datasets, displayed as an UpSet plot. Vertical bars represent the number of sites shared across datasets, while horizontal bars indicate the total number of sites per study. The sparse overlap highlights the experimental variability and condition-specific nature of the plant redoxome. (B) Maximum likelihood (LG+F+G) phylogenomic reconstruction of 37 representative Viridiplantae taxa. The tree spans the evolutionary trajectory from aquatic Chlorophyta to terrestrial Embryophyta, with distinct colors for each major taxonomic group. The bold backbone indicates the lineage leading to *Arabidopsis thaliana*, with internal nodes along this lineage indicated. All nodes along the path show high bootstrap support. Branch lengths represent substitutions per site (scale bar = 0.1).

To trace the evolutionary origins of these curated redox sites, a comprehensive phylogenomic backbone was constructed utilizing 37 representative *Viridiplantae* reference proteomes (Supplementary Table S1). The taxon sampling spanned the green lineage, ranging from aquatic Chlorophyta to Embryophyta. The resulting maximum-likelihood species tree (Figure 1B) recovered the consensus topology established by recent large-scale plant phylogenomics ^20^, ensuring a high-confidence scaffold for the analysis. To map the acquisition of redox-sensitive cysteines we labeled the inner nodes along the path from *Viridiplantae* root (N0) to Arabidopsis thaliana (N35). These nodes represent key evolutionary milestones, such as the transition to land (N16), the divergence of vascular plants (N20), spermatophytes (N23) and the emergence of flowering plants (N27).

Orthologous groups were identified using a reciprocal best BLAST hit (rBBH) strategy ^21^; ancestral sequence reconstruction (ASR) was then performed at each internal node to map the precise evolutionary node at which each cysteine residue was acquired. Cysteine gains along the evolutionary trajectory leading to *Arabidopsis* were classified into two distinct modes: **(1) Cys-protein co-gains**, where cysteine was present at the earliest inferred appearance of the gene, and **(2) Cys gain via amino acid replacement**, where cysteine arose later along the lineage leading to *Arabidopsis*. This approach aligned with previous evolutionary reconstructions of plant thiol-based switches ^22^. The final dataset comprised the gains mapping of 25,649 cysteine sites, distributed across 3,947 proteins onto ancestral nodes along the evolutionary trajectory leading to *A. thaliana*. Within this dataset, 709 RCys sites, 5,922 NRCys sites, and 19,018 UNCys sites were identified. Further breakdown of the 709 RCys sites revealed that 220 originated through amino acid replacement, while 489 resulted from Cys-protein co-gains. The relative frequency of gains along the evolutionary paths to Arabidopsis for each class is presented in Figure 2 alongside the rates of RCys gains. Because large-scale proteomic data inherently suffer from detection biases and stochastic peptide sampling limits, standard parametric statistical tests are frequently insufficient for evolutionary inference. Consequently, to rigorously test whether RCys gains were significantly enriched at specific nodes, permutation tests were performed with 10,000 iterations. In each iteration, the reactive label was randomly shuffled within two distinct pools: (1) the experimentally detected cysteines (RCys vs. NRCys), and (2) all cysteines within the sampled proteins (RCys vs. Total). This generated an empirical null distribution of expected RCys gains for each ancestral node in each pool.

**Figure 2.**
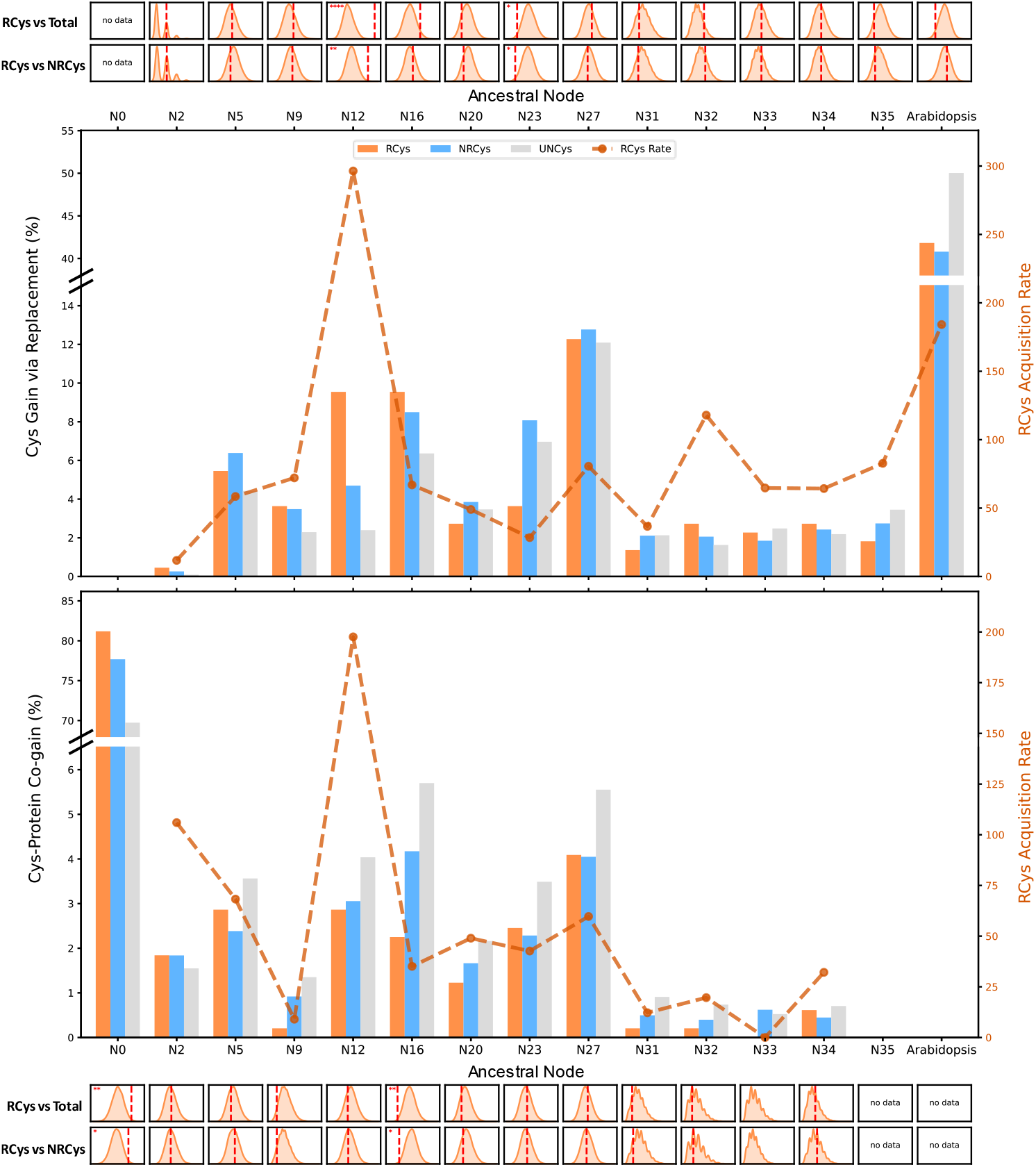
Evolutionary dynamics of cysteine gains and expansion of the redox proteome across the green lineage. Main panels display the relative frequency (left y-axis, bars) of cysteine sites acquired via amino acid replacement (Top) and Cys-protein co-gains conserved from the protein’s evolutionary root (Bottom). Cysteine sites are categorized as reactive (RCys, orange), non-reactive (NRCys, blue), and unclassified (UNCys, grey). The dashed orange line tracks the RCys Acquisition Rate (right y-axis), calculated by dividing the absolute number of acquired RCys sites at a given node by its preceding branch length. Left y-axes are broken to accommodate the high magnitude of gains at ancestral (N0) and terminal nodes. Histogram grids above and below the main charts display null distributions generated from 10,000 permutation iterations for the replacement and co-gain scenarios, respectively. Tests were conducted by randomly shuffling the “reactive” label between either the total protein cysteine pool (RCys vs Total) or the experimentally detected pool alone (RCys vs NRCys). Red dashed lines indicate the empirically observed RCys count at each node. Red asterisks denote statistically significant enrichment of observed RCys gains relative to the respective null models (* p < 0.05; ** p < 0.01; **** p < 0.0001). This temporal mapping highlights an ancient conserved redox core (N0) and a significant pre-terrestrial evolutionary burst (N12).

### Ancestral Core of the Redox Proteome

Notably, 397 RCys sites in 336 proteins, representing 81.18% of all Cys-protein co-gained reactive cysteines (489 in total), were inferred as present at the base of the *Viridiplantae* lineage (node N0). Importantly, comparing the gain frequencies of RCys, NRCys, and UNCys sites revealed a profound enrichment of RCys in Cys-protein co-gains at the tree root (Figure 2, bottom panel, permutation tests vs. NRCys, *p* = 0.0149; vs. total cysteines, *p* = 0.009). Functional enrichment analysis of the N0 co-gained RCys proteins revealed an enrichment of plastid-localized proteins (GO:0009532; p_adj = 1.74 × 10^−13^) and carbon fixation pathways (KEGG:00710; p_adj = 1.42 × 10^−6^), indicating that the foundational layer of the plant redox network, which includes regulation of photosynthesis, was already established in the common ancestor of *Viridiplantae*. The significantly higher frequency of RCys gains through protein co-gain at N0, compared with the expected baseline evolutionary signal (as estimated from comparable analyses of NRCys and UNCys sites), suggests that the gain of these regulatory sites was selectively favored, highlighting their functional importance.

Further examination of the evolutionary origin of RCys Cys-protein co-gains at the tree root found that 231 out of 336 (68.7%) of the proteins containing RCys had cyanobacterial homologs. Among these, 103 RCys, distributed across 94 proteins, were identified at the equivalent alignment position in at least one cyanobacterial proteome, suggesting that they represent ancient redox sensors conserved from the original cyanobacterial endosymbiont. Of these inherited sites, 47 (45.6%) were fully conserved across all four representative cyanobacterial proteomes, with a mean cysteine conservation of 81.3% across the panel. In contrast, the remaining 174 RCys sites (62.8%) showed no cysteine at the equivalent position in any cyanobacterial sequence, suggesting they emerged via amino acid substitutions during the evolutionary transition from the endosymbiotic event to the common ancestor of Viridiplantae (N0). This is consistent with the finding that primary plastid endosymbiosis coincided with a substantial expansion of the redox-sensitive proteome ^22^.

RCys within the chloroplastic ATP synthase gamma chain 1 (ATPC1) exemplify this deeply conserved ancestral core. Detailed evolutionary tracking of this highly conserved protein traces residues RCys139, RCys249, and UNCys255 back to the root of the green lineage at node N0 (Figure 3A). Notably, the cyanobacterial analysis revealed distinct evolutionary histories for these residues. More specifically, RCys139 was conserved across all four evaluated cyanobacterial proteomes, indicating that it is an ancient redox sensor directly inherited from the cyanobacterial endosymbiont. Conversely, the critical regulatory disulfide bridge formed by Cys249 and Cys255, whose thioredoxin-mediated reduction is essential for the light-dependent activation of the ATP synthase ^23^, was absent in cyanobacteria ^24,25^, indicating that this specific regulatory bond was not inherited, but rather emerged as an evolutionary adaptation in a pre-existing protein prior to the establishment of the green lineage at N0.

**Figure 3.**
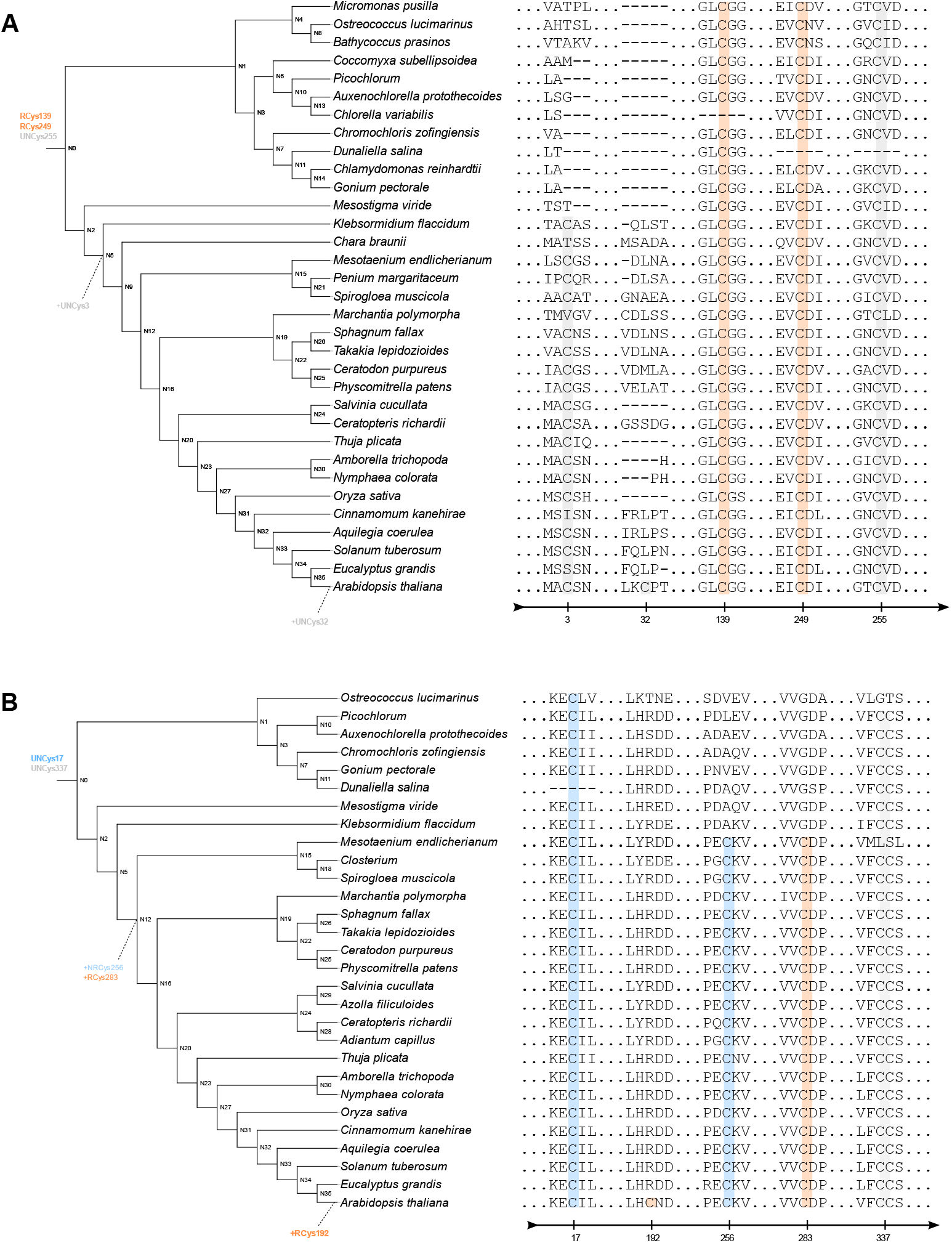
Chronological mapping of cysteine gains. Projected phylogenetic trees and an outline of the underlying multiple sequence alignments describe the Cys dynamics inferred from ancestral sequence reconstruction for two representative functionally distinct proteins. Ancestral nodes (N0–N35) denote key evolutionary divergence points. Cysteine residues are highlighted and color-coded based on their empirical reactivity labels in *Arabidopsis thaliana*: reactive (RCys, orange shading), non-reactive (NRCys, blue shading), and unclassified (UNCys, grey shading). Dotted arrows indicate the specific node where a novel cysteine was acquired via amino acid replacement. (A) ATPC1 (ATP synthase gamma chain 1): RCys139, RCys249 and UNCys255 are deeply conserved, and mapped to the ancestral root (Node N0) as Cys-protein co-gains. Subsequent non-regulatory replacements occurred later at N5 (+UNCys3) and the terminal *Arabidopsis* branch (+UNCys32). (B) GDI1 (Rab GDP-dissociation inhibitor 1): While the protein framework and ancestral cysteines (NRCys17, UNCys337) are conserved from N0, RCys283 and NRCys256 were acquired simultaneously via replacement exactly at the pre-terrestrial divergence (N12). A terminal replacement (+RCys192) is uniquely observed in the *Arabidopsis* branch.

### Pre-Terrestrial Expansion via Amino Acid Replacement

The ancestral core described above represents the inheritance of ancient aquatic redox machinery, already in place at the base of the green lineage. However, a fundamentally different pattern emerged when cysteine gains were traced through amino acid replacements in pre-existing proteins. This analysis revealed a pronounced burst of novel RCys gains at node N12, the evolutionary divergence point representing the common ancestor of *Zygnematophyceae* and *Embryophyta*. The frequency of RCys gains via amino acid replacement at this node spiked (Figure 2, top panel), and included 21 novel RCys regulatory switches, representing 9.8% of all RCys replacement gains. Permutation testing confirmed that this surge represents a statistically significant evolutionary burst against both background cysteine pools (vs NRCys, *p* = 0.0025; vs Total Cysteines, *p* = 0.001). Enrichment analysis of proteins acquiring novel RCys replacements at node N12 revealed significant representation in nucleotide metabolism (GO:0009117; *p_adj* = 4.3 × 10^−3^), Rab GDP-dissociation inhibitor activity (GO:0005093; *p_adj* = 1.1 × 10^−2^), and cell wall polysaccharide metabolism (GO:0010383; *p_adj* = 2.7 × 10^−2^). The disproportionate gain of novel thiol-based sensors at this node strongly suggests that an increasing proportion of proteins came under redox regulatory control in the algal ancestors of land plants. The significance of this pre-terrestrial expansion became even more apparent when evaluating the overall rate of evolutionary innovation. The RCys Acquisition Rate, which normalizes gains by the preceding evolutionary time (branch length), exhibited a massive, synchronized spike at node N12, with normalized RCys acquisition rates (co-gain: 197.58; amino acid replacement: 296.37). This rapid, synchronized accumulation of regulatory cysteines suggests that intense and diverse selection pressures drove the evolution of new redox-regulatory systems prior to terrestrialization.

A prime example of pre-terrestrial RCys gain via amino acid replacement is the Rab GDP-dissociation inhibitor 1 (GDI1) (Figure 3B). GDIs are key regulators of vesicular trafficking among endomembrane compartments ^26^, with GDI1 acting as an essential regulator of early embryogenesis in *Arabidopsis* ^27^. While GDI1 itself was consistently conserved since the root of the green lineage (node N0), its redox-regulatory capacity emerged much later via amino acid replacement. Specifically, RCys283 and NRCys256 were both acquired exactly at node N12. Furthermore, GDI1 independently acquired RCys192 in the terminal branch leading to *Arabidopsis*. This evolutionary gain may indicate that fundamental vesicular trafficking machinery was actively integrated into the redox-sensitive proteome just prior to terrestrialization.

### Post-Colonization Deceleration of Redox Innovation

The elevated rate of RCys acquisition observed at N12 was not sustained following the transition to land. At post-colonization nodes, the frequency of RCys gains fell significantly below permutation expectations in both gain scenarios; co-gains at the base of embryophytes (N16, vs NRCys *p* = 0.01; vs Total Cysteines *p* = 0.0086) and amino acid replacements at the divergence of seed plants (N23, vs NRCys *p* = 0.0163; vs Total Cysteines *p* = 0.0245) were each significantly depleted relative to the null distribution (Figure 2). This deceleration suggests that once the core terrestrial redox toolkit had been assembled, strong purifying selection constrained further large-scale remodeling of the redox proteome. This pattern, indicating the acquisition of reactive cysteines (RCs) at discrete evolutionary nodes rather than via a continuous, gradual process, may reflect an inherent trade-off between enhanced capacity for redox-sensitive regulation of activity and increased vulnerability to oxidative damage through overoxidation of RCs. Individual proteins nevertheless continued to acquire regulatory cysteines, as illustrated by the chloroplastic NAD(P)H-quinone oxidoreductase subunit N (NDHN; Figure 2S), which gained RCys188 at node N16 and NRCys122 at the base of angiosperms (N27). These gains indicate that post-colonization redox evolution proceeded through targeted, lineage-specific modifications rather than through a broad proteome-wide expansion that characterized the pre-terrestrial transition.

### Conclusion

The evolutionary mapping presented here suggests that the plant redox proteome evolved through an ancient establishment of a plastid-based metabolic core at the root of the green lineage, followed by a rapid, punctuated burst of regulatory innovation, achieved by systematically embedding novel thiol-based switches into pre-existing proteins, just before land colonization. Thus, the data presented here indicate that the capacity to fine-tune biochemical reactions via redox signaling represents a preadaptation that emerged in parallel with these structural and genomic adaptations.

It is important to note that this evolutionary framework carries an inherent interpretive limitation. The reactivity classifications (RCys, NRCys) used to label ancestral cysteine gains are defined exclusively by mass-spectrometric measurements in modern *Arabidopsis thaliana*. Cysteine reactivity, however, is an emergent biophysical property shaped by local electrostatics, pKa, and the surrounding residue microenvironment ^28^, none of which are directly recoverable from sequence reconstruction alone. Although current redox proteomics data are largely restricted to Arabidopsis, future studies in key algal lineages, including Zygnematophyceae representatives, combined with the development of AI-based approaches ^29^ for accurate prediction of cysteine reactivity, will enable a more comprehensive understanding of the major evolutionary events shaping contemporary redox proteomes.

## Author Contributions

R.O., E.H.C. and S.R. designed the research strategy. R.O. performed the computational analyses and data visualization. R.O., E.H.C. and S.R. wrote the manuscript. E.H.C. and S.R supervised the study. All authors reviewed and approved the final manuscript.

## Acknowledgments

We thank Yair Mau (Institute of Environmental Sciences, The Hebrew University of Jerusalem) for the valuable insights on the permutation analysis.

This research was supported by the European Research Council (ERC-COG, AGRIREDOX, grant no. 101086608) and the Israel Science Foundation (grant No. 1779/21).

## Resource Availability

Lead Contact: Further information and requests for resources and reagents should be directed to and will be addressed by the Lead Contact, Shilo Rosenwasser.

Materials Availability: This study did not generate new unique reagents or physical materials.

## Data and Code Availability

The raw liquid chromatography-tandem mass spectrometry (LC-MS/MS) datasets utilized to construct the cysteine reactivity map (Table S1) are publicly available through their original respective publications.

The 37 reference proteomes utilized for phylogenomic reconstructions (Table S1) are publicly accessible via the UniProtKB complete reference proteomes database, Phytozome v14, FernBase v1.2, and the NCBI RefSeq database.

All original code, custom Python parsing scripts, and configuration files generated for the reciprocal-best-BLAST-hit ortholog mapping and permutation testing are available from the lead contact upon request.

All additional information required to reanalyze the data reported in this paper is available from the lead contact upon request.

## Experimental Model and Study Participant Details

This study was purely computational and did not involve experimental models, organisms, or human/animal subjects.

## STAR Methods

### Proteomic Data Compilation and Reactivity Mapping

To compile the empirical dataset of redox-sensitive cysteines, peptide-level identifications were extracted from five independent large-scale *Arabidopsis thaliana* redox proteomics surveys. All extracted peptides were mapped against the *Arabidopsis thaliana* UniProt reference proteome (UP000006548). To standardize cross-study comparisons, residues were classified into three categories based on study-specific reactivity thresholds and fold-change cutoffs.

### Species Tree Reconstruction

To construct a comprehensive phylogenomic framework, 37 *Viridiplantae* reference proteomes were processed using OrthoFinder with the MAFFT option ^30,31^. This workflow was employed specifically to derive a concatenated species-level multiple sequence alignment (MSA) based on identified single-copy orthologs. A maximum-likelihood species tree was subsequently reconstructed from this concatenated MSA using IQ-TREE under the LG+F+G evolutionary model ^32^. Internal branch support was evaluated through 1,000 ultrafast bootstrap replicates, and the resulting phylogeny was manually rooted at the divergence between Streptophyta and Chlorophyta.

### Ortholog Identification and Gene Tree Alignment

High-confidence, one-to-one orthologs between the *A. thaliana* source genome and the 36 target species were established using a reciprocal-best-BLAST-hit (rBBH) pipeline executed in Python. The pipeline utilized NCBI BLAST+ v2.15.0 with an E-value threshold of 1e-5. Orthology was exclusively confirmed when the top-scoring forward search hit reciprocally identified the original query gene in the reverse search. Multiple sequence alignments (MSAs) for the resulting 21,485 ortholog groups were generated using MAFFT v7.505 ^31^ with default parameters. Gene trees were subsequently constructed by pruning the master species tree to include only the specific taxa present in each orthologous group, strictly preserving the established topological backbone.

### Ancestral Sequence Reconstruction and Evolutionary Mapping

Marginal ancestral protein sequences were reconstructed for all pruned gene trees utilizing FastML v3.11 ^33^. Reconstructions were performed using maximum-likelihood optimization (-ml) and gamma-distributed site rates (-qf). Amino acid gain events were systematically called by scanning the reconstructed ancestral sequences iteratively from the *A. thaliana* terminal leaf back to the root. An evolutionary gain was recorded when the state at an aligned amino acid position changed between a parent and child node, provided that the marginal probability of the ancestral site exceeded a confidence threshold of 0.90. Identified gains were subsequently classified as “Cys-protein co-gain” when the residue was present at the deepest inferred node of the specific protein or “amino acid replacement” when the residue arose via amino acid replacement.

### Identification of Cyanobacterial Homologs and Residue-Level Cysteine Conservation Analysis

To determine the prokaryotic origins of the ancestral redox core, Arabidopsis thaliana protein sequences that acquired reactive cysteines (RCys) at the root of the green lineage (Node N0) were screened for homology against a panel of four representative cyanobacterial proteomes (Table S1). Homology searches were performed using PSI-BLAST (NCBI BLAST+ v2.15.0) with five iterations, an E-value threshold of 1e-3, a PSSM inclusion threshold of 1e-2, 500 maximum target sequences per iteration, and the BLOSUM45 substitution matrix, which is suited to the ∼20–35% sequence identity expected at this evolutionary distance. Residue-level conservation analysis was performed by aligning each query with its cyanobacterial homologs using MAFFT v7.505 (L-INS-i). For each RCys site, the ungapped residue position was mapped to its corresponding multiple sequence alignment column and inspected for cysteine presence across all cyanobacterial homologs. Conversely, RCys sites present at Node N0 but absent across the entire cyanobacterial panel were classified as those which emerged via amino acid substitutions during the evolutionary transition from the primary endosymbiotic event to the common ancestor of Viridiplantae.

### Permutation Testing for Node Enrichment

An empirical null distribution was generated using a protein-matched permutation approach, to assess whether the evolutionary gain of RCys occurred more frequently than expected by chance. For each protein containing n observed RCys, an equivalent protein was randomly sampled from the reactivity map that possessed enough total cysteines to accommodate the labels. The RCys labels were then randomly shuffled among the available cysteine residues within the sampled proteins. This process was conducted independently for two distinct background pools: a detected pool restricted to experimentally identified residues (RCys + NRCys) and a “total” pool encompassing all cysteines within the detected proteins (RCys + NRCys + UNCys). After reassigning labels, the randomized reactive sites were mapped to their inferred evolutionary gain nodes, and the total number of RCys gains per node was recalculated across the phylogeny. This procedure was repeated for 10,000 permutations to generate a robust null distribution. Empirical p-values were calculated as the proportion of permutations yielding RCys gain counts equal to or greater than the true observed counts, with significance defined as p < 0.05.

### Functional Enrichment Analysis

Functional enrichment analyses were conducted utilizing the g:Profiler web server ^34^. Target gene lists were rigorously composed of *A. thaliana* orthologs that definitively acquired RCys at specific nodes of interest and specific gain scenarios (N0 co-gains and N12 substitution gains). The statistical background pool was strictly restricted to the list of all *A. thaliana* proteins detected in the integrated proteomics datasets, ensuring that enrichment results were not systemically biased by highly abundant or easily ionized proteins. Enrichment across Gene Ontology classifications (Biological Process, Molecular Function, Cellular Component) and KEGG pathways was tested using Fisher’s exact test, with statistical significance determined after Benjamini-Hochberg False Discovery Rate (FDR) correction at an adjusted p-value threshold of ≤ 0.05.

