## Supplementary Figures for "Expansion of Plant Redox Protein Network Predated Plant Terrestrialization"

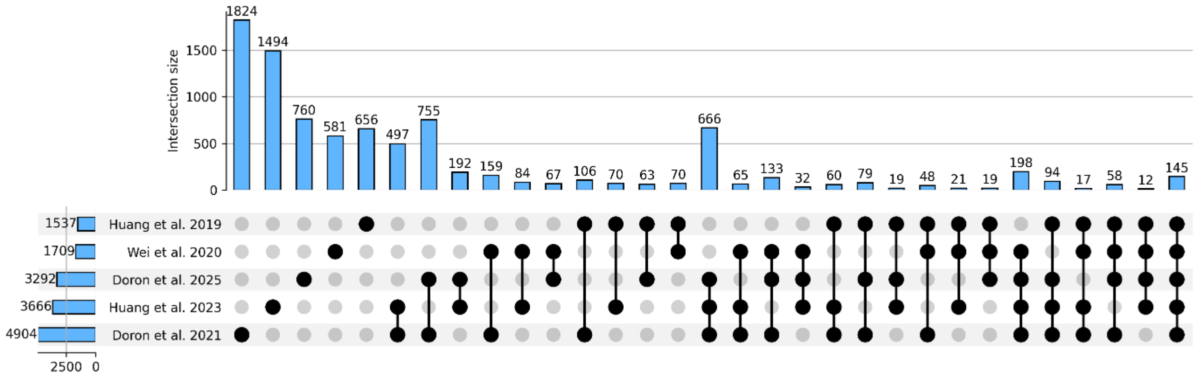


**Figure S1. Overlap of total cysteine sites across five independent Arabidopsis thaliana redox proteomics datasets, related to Figure 1.**

UpSet plot displaying the intersection of all detected cysteine sites (RCys, NRCys, and UNCys) across the five integrated studies: $H_2O_2$-induced oxidation in whole-cell lysates (Huang et al., 2019), disulfide-linked peptide reporters in cell cultures (Wei et al., 2020), intact leaf profiling (Doron et al., 2021), excess light stress mapping (Huang et al., 2023), and dark-to-light photosynthetic induction (Doron et al., 2025). Vertical bars represent the size of each intersection (shared sites between specific study combinations), while horizontal bars indicate the total number of cysteine sites per study. Out of 4,904 total detected sites in Doron et al. 2021, the largest single study, only 666 sites were consistently shared among the three largest datasets. This sparse overlap underscores the complementary nature of the methodologies employed and the condition-specific detection of cysteine sites, reinforcing the rationale for integrating multiple independent datasets to achieve comprehensive coverage of the Arabidopsis redoxome.

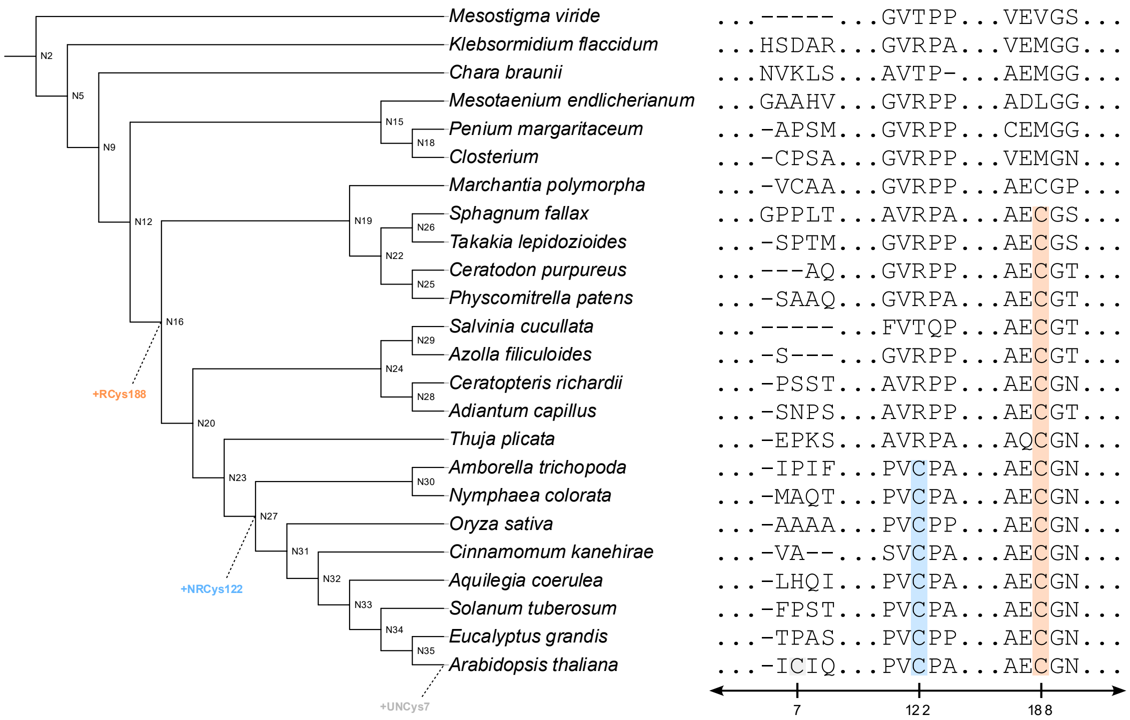


**Figure S2. Evolutionary tracking and post-colonization fine-tuning of the chloroplastic NAD(P)H-quinone oxidoreductase subunit N (NDHN), related to Figure 3.**

Projected phylogenetic tree and multiple sequence alignment (MSA) illustrating the cysteine dynamics of the NAD(P)H-quinone oxidoreductase subunit N (NDHN) as inferred from ancestral sequence reconstruction. NDHN is an oxygenic photosynthesis-specific subunit that provides structural stability to the NDH-1 complex by interfacing with NdhV; its functional necessity is demonstrated by the inability of *ndhN* mutants to survive under ambient $CO_2$ concentrations. The ancestral protein emerged at Node N2. As land plants evolved, the protein sequentially accrued regulatory complexity, acquiring the reactive switch RCys188 at the common ancestor of Embryophyta (Node N16), followed by a non-reactive cysteine (NRCys122) at the divergence of flowering plants (Node N27), and a terminal UNCys7 addition uniquely within the *Arabidopsis thaliana* lineage. Ancestral nodes (N0–N35) represent key evolutionary divergence points. Cysteine residues are color-coded based on empirical reactivity labels from *A. thaliana* datasets: reactive (RCys, orange shading), non-reactive (NRCys, blue shading), and unclassified (UNCys, grey shading). Dotted arrows highlight the specific nodes where novel cysteines were acquired through amino acid replacement.
